# The effect of APOE genotype and streamline density volume, on hippocampal CA1 down-regulation: a real-time fMRI virtual reality neurofeedback study

**DOI:** 10.1101/643577

**Authors:** Stavros Skouras, Jordi Torner, Patrik Anderson, Yury Koush, Carles Falcon, Carolina Minguillon, Karine Fauria, Francesc Alpiste, Juan D. Gispert, José L. Molinuevo, the ALFA Study

**Affiliations:** Barcelonaβeta Brain Research Center (BBRC), Pasqual Maragall Foundation, Barcelona, Spain; School of Engineering and Design, Polytechnic University of Catalonia, Barcelona, Spain; Centro de Investigación Biomédica en Red de Bioingeniería, Biomateriales y Nanomedicina (CIBER-BBN), Madrid, Spain; SUBIC, Stockholm University, Stockholm, Sweden; Department of Radiology and Biomedical Imaging, Yale University; CIBER Fragilidad y Envejecimiento Saludable (CIBERFES), Madrid, Spain; Universitat Pompau Fabra, Barcelona, Spain

## Abstract

Hippocampal hyperactivity is a precursor of Alzheimer’s disease and more prominent in APOE-ε4 carriers. It is therefore important to investigate the processes of hippocampal self-regulation, to monitor therapeutic efficacy of preclinical interventions. We have developed a closed-loop, virtual reality neurofeedback paradigm for real-time fMRI, that provides a standardized method for quantifying processes of hippocampal self-regulation. We acquired multi-modal neuroimaging data from a sample of 53 cognitively unimpaired subjects at risk for AD and applied standard methods of structural and functional connectomics. The analyses reveal significant negative associations between hippocampal CA1 down-regulation performance and APOE-ε4 alleles, as well as hippocampal streamline density volume. Better memory performance was associated with increased, bilateral hippocampal functional connectivity during the neurofeedback task. These are the first results to link neurofeedback performance to a genetic risk factor and structural connectivity. Further, these are the first evidence that functional cohesion between the hippocampi can reflect subtle differences in memory function, in cognitively unimpaired individuals at risk for AD. We provide a novel method to assess hippocampal function in preclinical AD, and propose it can be used to derive proxies for neural reserve.

**Highlights:**

- APOE-ε4 alleles impact hippocampal down-regulation neurofeedback performance.
- Hippocampal streamline density volume is associated with decreased hippocampal down-regulation performance.
- Bilaterally cohesive hippocampal activity is associated with better memory performance.
- We provide a novel paradigm to investigate self-regulation and brain function.

## Introduction

Recent fMRI studies have renewed hope for identifying subtle functional changes preceding irreversible neurodegeneration due to Alzheimer’s disease (AD) [1–3]. In-depth literature review suggests that the hippocampus is the only brain region that is consistently and bilaterally both a) implicated in memory encoding processes, as indicated by a meta-analysis of 72 fMRI studies of subsequent memory effects [4] and b) differing in activity between AD patients and controls, as indicated by a quantitative meta-analysis of 11 fMRI studies on memory encoding [5]. Moreover, recent evidence suggests hippocampal hyperactivity as a precursor of amyloid deposition [1], which is accepted as the earliest sign of preclinical AD [6]. Similarly, APOE-ε4 carriers, exhibit hippocampal hyperactivity during a memory task designed to assess pattern separation [7], suggesting the possibility of a genetic mechanism linked to the concepts of neural reserve and neural efficiency [8–9]. Therefore, it is important to investigate the processes of hippocampal self-regulation, to improve detection of the earliest stages of the pathophysiological continuum of AD and to monitor therapeutic efficacy of clinical interventions.

White matter microstructure is altered in cognitively normal middle-aged APOE-ε4 homozygotes [10] and a previously established increase in default mode network (DMN) connectivity during β-amyloid accumulation [2] appears to be compensating for decreasing functional connectivity of the left hippocampus [3], that specializes in the representation of objects’ episodic context [11]. Age and psychiatric pathology exert significant effects on neurofeedback (NF) performance of the DMN [12] and we have developed an open-source virtual reality (VR) NF environment for real-time fMRI (rt-fMRI), to systematize further investigations [13].

In previous studies of healthy young adults, rt-fMRI NF performance correlated with down-regulation, but not up-regulation, of the parahippocampal formation [14]. Moreover, learning to up-regulate the left dorsolateral prefrontal cortex was associated with higher improvement of working memory, across two sessions [15]. Virtual reality has also shown promise for the development of immersive experimental tasks with increased ecological validity [16], that have revealed significant insights regarding hippocampal function [11, 17–18]. A crucial consideration in combining VR and NF towards translational applications, regards the accurate selection of purpose-specific NF target regions [19]. Advanced neuroimaging methods have enabled the probabilistic segmentation of hippocampal subfields from T1-weighted anatomical images [20–23]. Evidence suggests the dorsal hippocampal subfield cornu ammonis 1 (CA1) to be the part of the hippocampus that selectively experiences atrophy due to AD compared to normal aging [22–23].

We reasoned that if preclinical alterations in brain function occur prior to neurodegeneration, it is probable that they are most prominent in areas that later undergo atrophy due to AD. Using a VR NF paradigm that was custom-made for this study [13], we measured the ability to exert voluntary control on the activity of hippocampal subfield CA1, in subjects at risk for AD. We hypothesized that memory performance, structural connectivity and the APOE genotype, would exert distinct effects on hippocampal down-regulation NF performance.

## Methods

### Participants

Participants comprised of 53 adult volunteers (age M = 62.77 years; SD = 5.73) from the ALFA (Alzheimer’s and Families) cohort, that features 86.3% descendants of AD patients and 19% APOE-ε4 carriers, hence presenting increased risk for AD [24]. All participants had no neurological or psychiatric history and at the time of scanning were cognitively normal, according to their scores on the retention index (M = 0.98; SD = 0.06) of the Free and Cued Selective Reminding Test (FCSRT) [25], that had been measured within the preceding six months. Participants had previously also completed the Cognitive Reserve Questionnaire (CRQ), a questionnaire comprised of 10 questions whose total score serves as a proxy for cognitive reserve [26] (M = 16.34; SD = 3.91). Thorough quality control was applied in advance, to exclude datasets having more than 10% invalid functional volumes due to movement and acquisitions during which technological complications, fatigue, discomfort or sleep had occurred. The local ethics committee ‘CEIC-Parc de Salut Mar’ reviewed and approved the experimental protocol, that was in accordance with the Declaration of Helsinki.

### Apolipoprotein E genotyping

Participants’ APOE genotype was determined as previously described [24]. Briefly, proteinase K digestion followed by alcohol precipitation, was used to obtain DNA from the blood cellular fraction. Samples were genotyped for two Single Nucleotide Polymorphisms (SNPs), rs429358 and rs7412, so that the number of APOE-ε4 alleles could be determined for each participant. Results are displayed in Table 1.

**TABLE 1:** Demographics and descriptive statistics.

| | Entire sample | APOE-$\epsilon 4$<br>non-carriers | APOE-$\epsilon 4$<br>heterozygotes | APOE-$\epsilon 4$<br>homozygotes | APOE-$\epsilon 4$<br>group effect |
| --- | --- | --- | --- | --- | --- |
| | $N = 53$ | $n_a = 14$ | $n_b = 29$ | $n_c = 10$ | |
| Sex ratio<br>(M / F) | 21 / 32 | 4 / 10 | 10 / 19 | 7 / 3 | $\chi^2(2, N=53) = 4.89$<br>$P = 0.087$ |
| **Age** | M = 62.786<br>SD = 5.731 | M = 65.408<br>SD = 1.552 | M = 63.962<br>SD = 5.752 | M = 55.705<br>SD = 3.540 | F(2,50) = 14.90<br>P < 0.001 |
| **FCSRT**<br>**Total recall** | M = 44.340<br>SD = 3.695 | M = 43.000<br>SD = 5.218 | M = 44.138<br>SD = 2.973 | M = 46.800<br>SD = 1.549 | F(2,50) = 3.48<br>P = 0.038 |
| **FCSRT**<br>**Retention Index** | M = 0.983<br>SD = 0.060 | M = 0.984<br>SD = 0.079 | M = 0.987<br>SD = 0.057 | M = 0.969<br>SD = 0.044 | F(2,50) = 0.33<br>P = 0.722 |
| **CRQ score** | M = 16.340<br>SD = 3.912 | M = 15.929<br>SD = 4.305 | M = 16.448<br>SD = 3.766 | M = 16.600<br>SD = 4.142 | F(2,50) = 0.11<br>P = 0.899 |
| **Hippocampal**<br>**volume / TIV** | M = 0.005<br>SD = 0.001 | M = 0.005<br>SD = 0.001 | M = 0.0053<br>SD = 0.001 | M = 0.0052<br>SD = 0.000 | F(2,50) = 0.58<br>P = 0.564 |
| **Hippocampal**<br>**SVD** | M = 0.156<br>SD = 0.051 | M = 0.170<br>SD = 0.073 | M = 0.154<br>SD = 0.045 | M = 0.146<br>SD = 0.020 | F(2,50) = 0.72<br>P = 0.492 |
| **Hippocampal**<br>**down-regulation**<br>**NF score** | M = -0.041<br>SD = 0.4873 | M = 0.125<br>SD = 0.538 | M = -0.029<br>SD = 0.492 | M = -0.311<br>SD = 0.274 | F(2,50) = 2.49<br>P = 0.093 |

*APOE-ε4 homozygotes were younger and performed better in total recall but not in retention. For all subsequent statistical tests neurofeedback performance and total recall were orthogonalized to the APOE genotype, as well as sex, age, CRQ score and hippocampal volume. Abbreviations: M ~ male; F ~ female; FSCRT ~ free and cued selective reminding test; CRQ ~ cognitive reserve questionnaire; TIV ~ total intracranial volume; SDV ~ streamline density volume; NF ~ neurofeedback.*

### Image acquisition

All scanning was performed in a single 3 T Philips Ingenia CX MRI scanner (2015 model). Prior to the real-time NF session, scanning comprised of a 3D T1-weighted image with the following parameters: Repetition Time (TR) = 9.90 ms, Echo Time (TE) = 4.60 ms, Flip Angle = 8, voxel resolution of 0.75×0.75×0.75 mm3 on 240 sagittal slices; and a diffusion-weighted sequence with parameters: TR = 9000 ms, TE = 90 ms, Flip Angle = 90, 72 non-collinear directions (b = 1300 s/mm2) and 1 nongradient volume (b = 0 s/mm2), voxel size of 2.05×2.05×2.20 mm2 on 66 axial slices. During rt-fMRI, echo planar imaging was used with an echo time of 35 ms and a repetition time (TR) of 3 s. Slice acquisition was interleaved within the TR interval. The matrix acquired was 80×80 voxels with a field of view of 240 mm, resulting in an in-plane resolution of 3 mm. Slice thickness was 3 mm with an interslice gap of 0.2 mm (45 slices, whole brain coverage).

### Structural image processing

T1-weighted images were subjected to the standard procedures implemented in the voxel-based morphometry toolbox (VBM toolbox; RRID:SCR_014196), to identify potential outliers (there were no outliers). T1 images were independently subjected to the standard FreeSurfer pipeline. The Connectomizer (v2.4.1; by Marcel A. de Reus; Dutch Connectome Lab, UMC Utrecht) was used to identify streamlines between all possible pairs of regions of interest (ROIs), based on the Cammoun Desikan-Killiany parcellation maps [27] (for extensive details please refer to [28]). With Matlab (version 9.3; Mathworks RRID:SCR_001622), the brain regions that featured consistent bilateral streamlines to the hippocampi across all participants were identified. These regions were the entorhinal cortex, amygdala, thalamus, posterior cingulate and parahippocampal formation. Using these regions only, the average hippocampal streamline density volume (SDV) [29–30], which describes the number of streamlines between two ROIs normalized by the sum of their volumes, was computed for each participant. FreeSurfer output was also used to compute each participant’s normalized hippocampal brain volume, as a proxy of brain reserve, by dividing the sum of the volume of the right and left hippocampi by each participant’s total intracranial volume [31]. The N4 nonparametric non-uniform intensity normalization bias correction function [32–33] of the Advanced Normalization Tools [34] (ANTs v2.x; RRID: SCR_004757) was applied on all T1 images, followed by an optimized blockwise non-local means denoising filter [35]. Multi-atlas segmentation with joint label fusion [36] was used to segment each participant’s bilateral hippocampal subfields, and derive probabilistic maps for CA1. These maps were thresholded at P = 0.9 to create binary masks that were used as NF target ROIs.

### VR task and real-time NF

The VR environment and real-time computations have been described in detail elsewhere [13]. Briefly, participants were immersed in a VR environment developed in the game-engine Unity (Unity Technologies ApS, San Francisco, CA, US), that resembles a park, with trees, a lake, a bridge etc. (Fig. 1). During a 30-minute-long scanning session, participants could run inside the VR environment in a fixed, circular path. Their task was to explore different mental strategies with the aim of achieving the maximum velocity and traversing the maximum distance, while attending to the VR environment and trying to remember its features. Following the first 30 functional volumes, that served to establish a baseline, with every TR, the most recent change in hippocampal CA1 activity was compared to normative measures of change derived from the preceding 30 scans. Increases in hippocampal activity triggered a 5% decrease of velocity and decreases in hippocampal activity triggered a 5% increase of velocity in the VR environment. Every 90 seconds, the velocity was reset to 50%. Apart from the visual experience of movement, the current velocity was displayed as a percentage of the maximum possible velocity and a green or red signal, superimposed on a coronal brain view, reflected the direction of the most recent change (Fig. 1). This setup encapsulated principles of operant conditioning, reinforcement learning and sensory-aided learning, encouraging improvement of performance even passively, through positive valence associated with acceleration and negative valence associated with deceleration. The purpose of the task was to drive CA1 activity in each participant to the minimum possible. Real-time image preprocessing comprised of rigid-body registration to an initial reference volume, temporal high-pass filtering at a cut-off frequency of 1/200 Hz [37], and voxel efficiency weighting [19] through voxel-wise normalization within the 30-volume sliding time-window (for extensive details please refer to [13]).

**Fig. 1:**
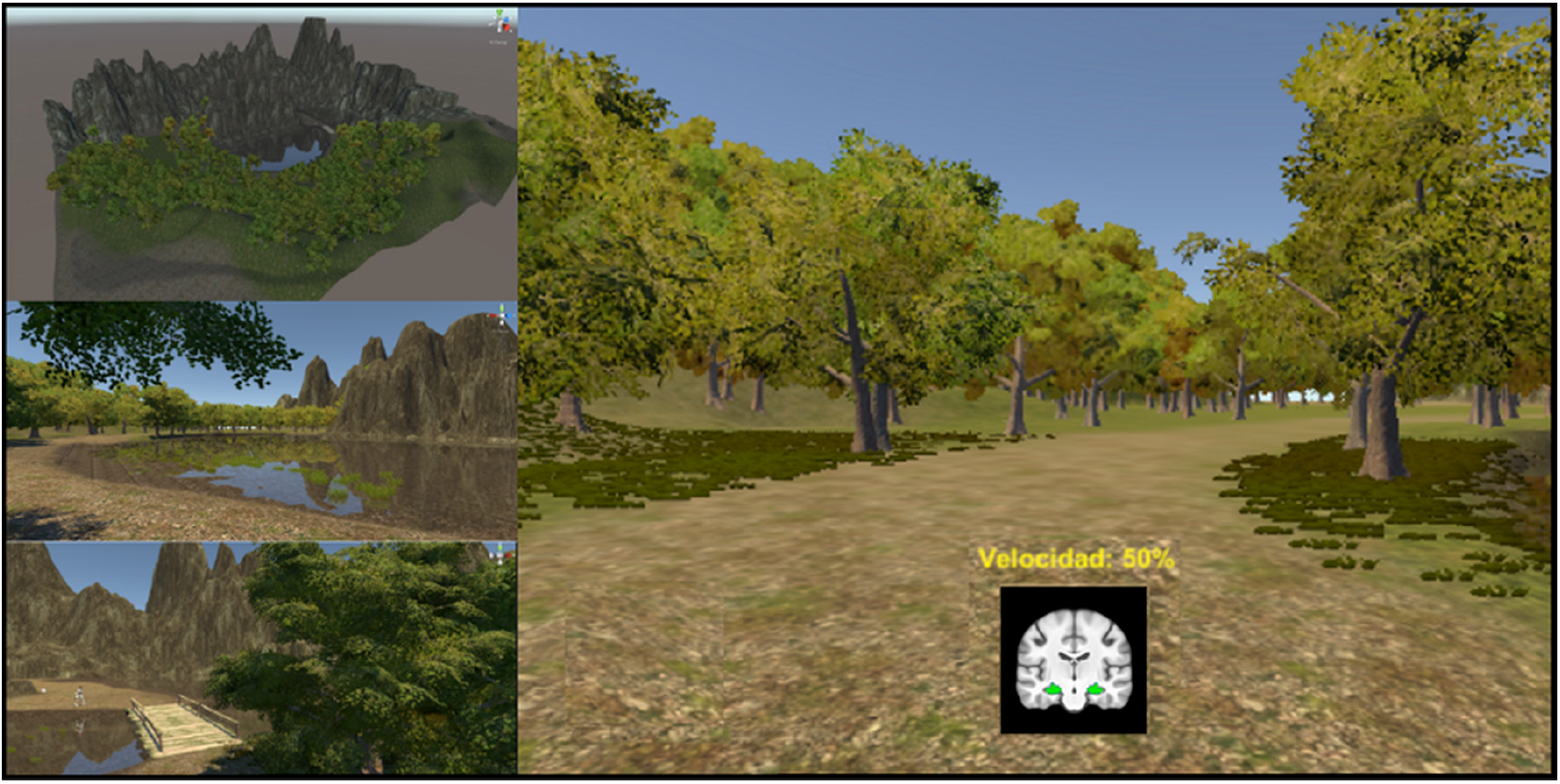
Virtual reality environment. Graphical snapshots illustrate the immersive virtual reality environment developed for the study. The velocity was updated online with every new functional volume, resulting in 570 NF signals across the session. Through sensory-aided operant conditioning, pleasant accelerations and unpleasant decelerations acted as positive and negative reinforcers respectively. Adaptive task difficulty ensured a standardized level of visual stimulation and similar overall experience for all participants, thereby controlling for important perceptual confounds.

### Functional image processing

Offline functional image preprocessing comprised of standard procedures using SPM12 (Statistical Parametric Mapping, RRID:SCR_007037) and the MIT connectivity toolbox (Connectivity Toolbox, RRID:SCR_009550), featuring slice-time correction, realignment and reslicing of functional volumes, denoising via regression of average white matter timeseries, average cerebrospinal fluid (CSF) timeseries, 24 Volterra expansion movement parameters and scan-nulling regressors [38] produced by the Artifact Detection Tools (ART; RRID: SCR_005994). Functional data and CA1 ROIs were normalized to the ICBM MNI template, which features sharp resolution, detailed gyrification and high signal-to-noise ratio, with minimum bias [39], via a custom template derived from the same population [3], using SyGN [33, 40]. Finally, using the Leipzig Image Processing and Statistical Inference Algorithms (LIPSIA v2.2.7 (2011), RRID:SCR_009595; [41]), a temporal highpass filter with a cut-off frequency of 1/90 Hz was applied to remove low frequency drifts in the fMRI time series, and spatial smoothing was performed using a 3D Gaussian kernel and a filter size of 6 mm at FWHM. Functional connectivity analysis was performed using the average of each subject’s normalized bilateral CA1 as the seed timeseries.

### Statistical Analysis

First-level GLM was employed to quantify NF performance per participant. Using Pearson’s r as a metric of similarity, actual NF moment-to-moment regulation, as captured in the realigned average CA1 timeseries, was compared to a linear vector representing the target performance of continuous down-regulation, producing a measure of NF performance for each participant. First, we investigated whether the unique variance associated with episodic memory was reflected in the functional connectivity of hippocampal CA1 during the NF task. Total Recall (TR), which constitutes the signature of memory impairment that is specific to AD [42–43], was orthogonalised to potential confounding variables; specifically sex, age, APOE genotype, hippocampal volume, cognitive reserve and NF performance. This resulted in an orthogonal 2^nd^-level design matrix that was used for GLM of the unique effect of TR on CA1 functional connectivity during VR NF. Whole-brain functional connectivity maps were corrected for multiple comparisons using 10,000 iterations of Monte Carlo simulations, with a pre-threshold of Z > 2 and a corrected significance level of P < 0.05. We also assessed the direct effect of TR and the FCSRT Retention Index on NF performance, corrected for sex, age, APOE genotype, hippocampal volume and cognitive reserve. Similarly, we investigated for a possible correlation between NF performance and hippocampal volume or cognitive reserve. We used non-parametric correlation testing to investigate the effect of the number of APOE-ε4 alleles on NF performance, corrected for sex, age, hippocampal volume and cognitive reserve. Last, we used correlation testing to investigate the effect of SDV on NF performance, corrected for sex, age, hippocampal volume and cognitive reserve. For all orthogonalisations we used the recursive Gram-Schmidt orthogonalisation implemented in SPM12 (SPM, RRID:SCR_007037), following standardization of scale variables. For each statistical test, we ensured that the variable of interest was completely orthogonal to all covariates in the design matrix, as well as to all original vector data, while at the same time remaining in high correlation with its own original vector. This was done to ensure that we investigated effects due to the unique variance in the variable of interest, without deviating from meaningful observations.

### Code and Data accessibility

The VR environment is available as open-source software via the ‘VR_multipurpose_v1.0’ repository (GitHub; RRID:SCR_002630). Data used and software developed for the analysis can been made available to researchers for non-commercial purposes, following agreement and approval by the Barcelonabeta Brain Research Center’s Data and Publications Committee.

## Results

### Relation of functional connectivity and NF performance with Memory performance

Controlling for age, sex, cognitive reserve, APOE genotype, hippocampal volume and NF performance, TR correlated significantly with increased functional connectivity between CA1 and the dorsal hippocampus bilaterally (Table 2). That is, participants that performed better in the memory test, also modulated activity in dorsal CA1 more cohesively across the two hemispheres, during the VR neurofeedback task. Controlling for age, sex, cognitive reserve, APOE genotype and hippocampal volume, there was a non-significant trend for negative correlation between NF performance and the FCSRT measures of TR [r = −0.045, p = 0.746] and Retention Index [r = −0.263, p = 0.057].

**TABLE 2.** Correlation of FCSRT TR with whole-brain, voxel-wise FC maps (seed: bilateral mean CA1), corrected for whole-brain multiple comparisons (P < 0.05)

| Region | MNI coordinates | Cluster size (mm<sup>3</sup>) | z-value |
| --- | --- | --- | --- |

|  |  |  | max (mean) |
| --- | --- | --- | --- |
| Right dorsal CA1/CA2 | 33 -21 -11 | 594 | 3.73 (2.89) |
| Left dorsal CA1/CA2 | -36 -27 -11 | 297 | 3.85 (2.90) |

*The outermost right column indicates the maximal z-value of voxels within a cluster (with the mean z-value of all voxels within a cluster in parentheses). Abbreviations: FCSRT ~ Free and Cued Selective Reminding Test; FC ~ Functional Connectivity; MNI ~ Montreal Neurological Institute; CA1 ~ Cornu Ammonis 1; CA2 ~ Cornu Ammonis 2.*

### Influence of the APOE genotype on NF performance

Controlling for sex, age, hippocampal volume, cognitive reserve and TR, NF performance showed a significant negative correlation with the presence of APOE-ε4 alleles in a dose-dependent manner, r = −0.288, p = 0.037; Fig. 4.

**Fig. 2.**
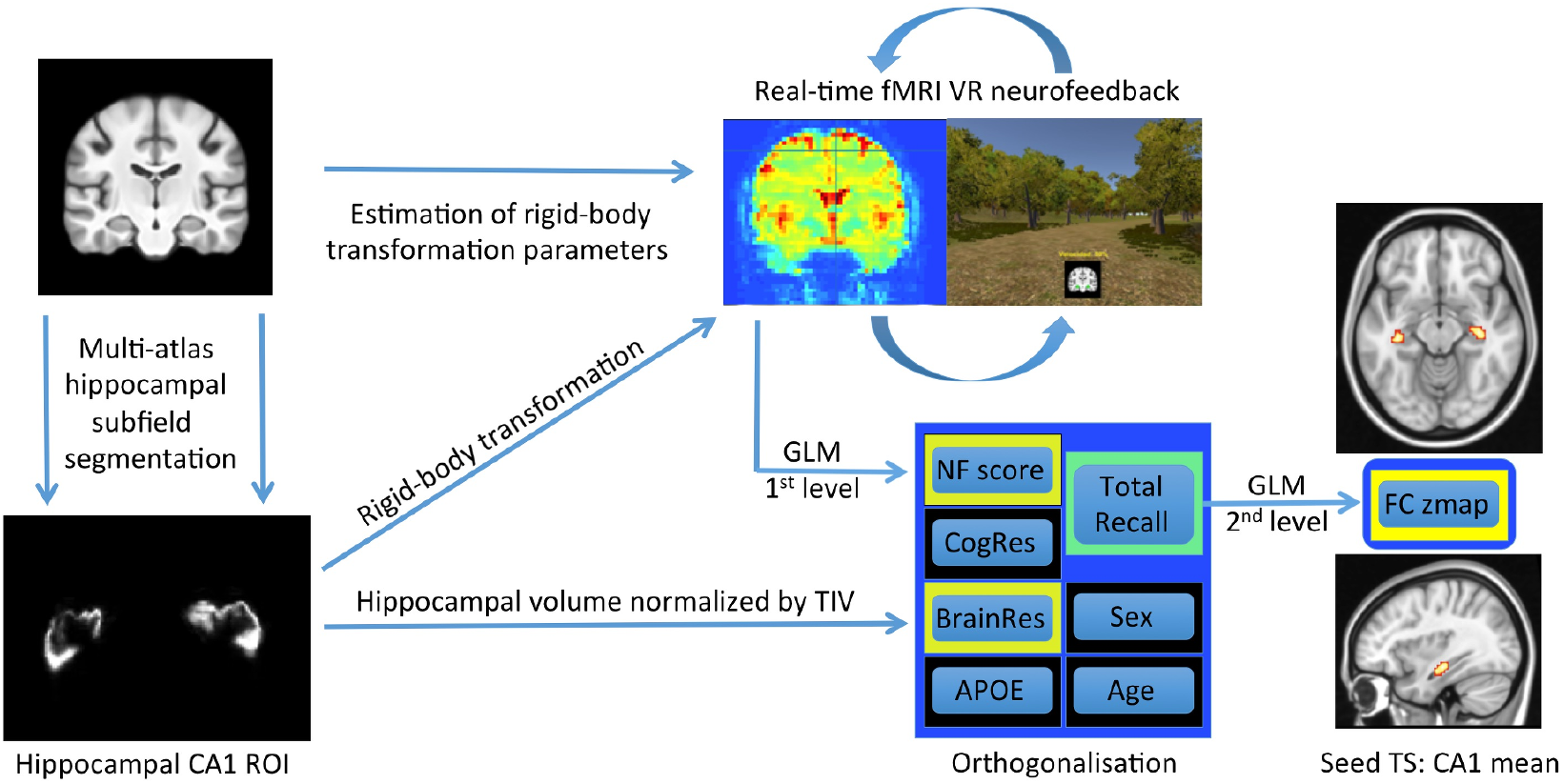
Illustration of the neuroimaging pipeline. Prior to the NF session, using previously acquired anatomical images, multi-atlas hippocampal subfield segmentation localized and segmented hippocampal subfield CA1. At the beginning of the NF session, rigid-body transformation was used to register the ROI mask to the rt-fMRI data. During the NF session, moment-to-moment changes in hippocampal CA1 activation were estimated via voxel efficiency weighting and reference activity from a sliding window of 30 volumes. Changes in activation effected inverse changes of velocity in the VR environment with every new functional volume, resulting in 570 NF signals. Following the NF session, offline GLM implementing regression against a target vector was used to derive a conventional measure of NF regulation performance. Additionally, by providing 570 volumes of uninterrupted hippocampal regulation, the paradigm was optimized for computing the functional connectivity of the region of interest (i.e. identifying the brain regions that correlate with hippocampal activity during NF). Abbreviations: VR: virtual reality; CA1: cornu ammonis 1; TIV ~ total intracranial volume; GLM: general linear model; NF: neurofeedback; CogRes: cognitive reserve; BrainRes: brain reserve; APOE: apolipoprotein genotype; FC: functional connectivity; TS: timeseries.

**Figure 3.**
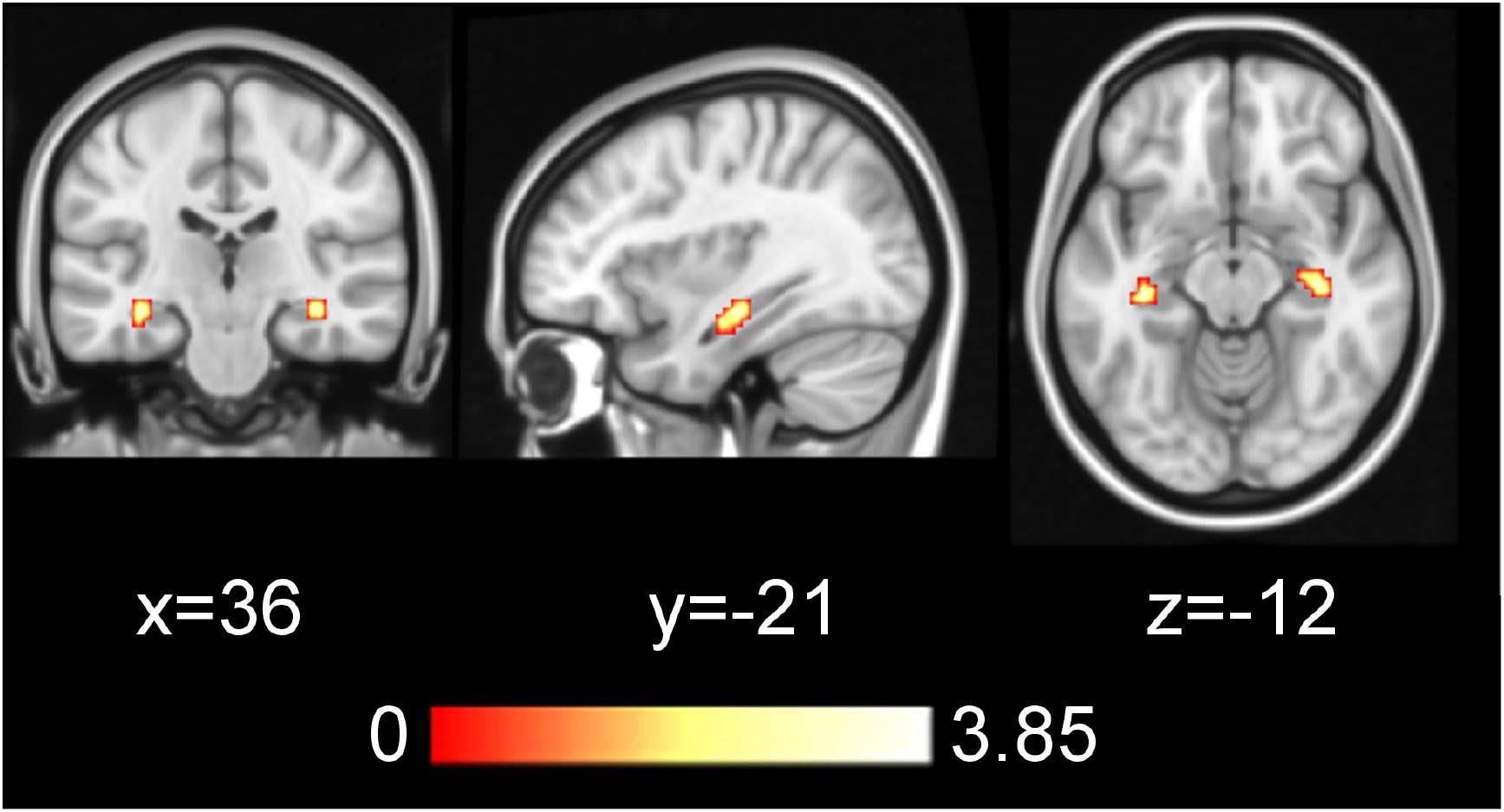
**Positive correlation between Total Recall (FCSRT) and mean CA1 functional connectivity**, during VR NF, corrected for sex, age, APOE genotype, hippocampal volume, cognitive reserve and neurofeedback performance (z > 2, P < 0.05 whole-brain corrected). This finding suggests that during the VR NF task, dorsal CA1 activity shows more cohesion between the two hemispheres in participants with good memory performance. ROI masks of these results are available for future investigations (see section Code and Data Availability).

**Figure 4.**
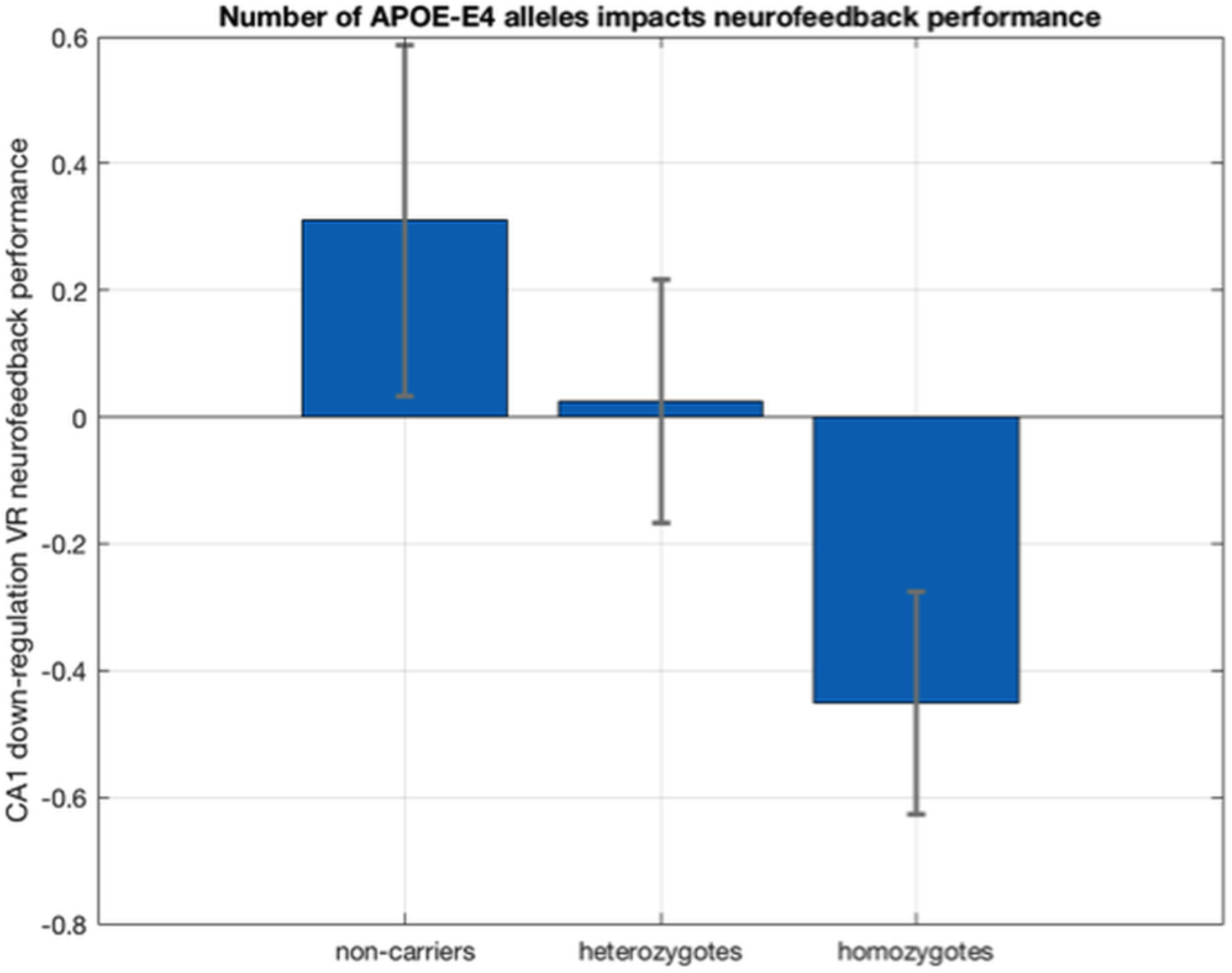
Dose-dependent effect of APOE-ε4 alleles on CA1 down-regulation performance. Hippocampal CA1 down-regulation performance scores were corrected for sex, age, hippocampal volume, cognitive reserve and memory performance [r = −0.288, p = 0.037].

### Relation of structural connectivity with NF performance

Controlling for sex, age, APOE genotype, hippocampal volume, cognitive reserve and TR, NF performance correlated with left mean hippocampal SDV, r = −0.371, p = 0.006 (Fig. 5). This was not the case for the right hippocampus r = −0.101, p = 0.469.

**Figure 5:**
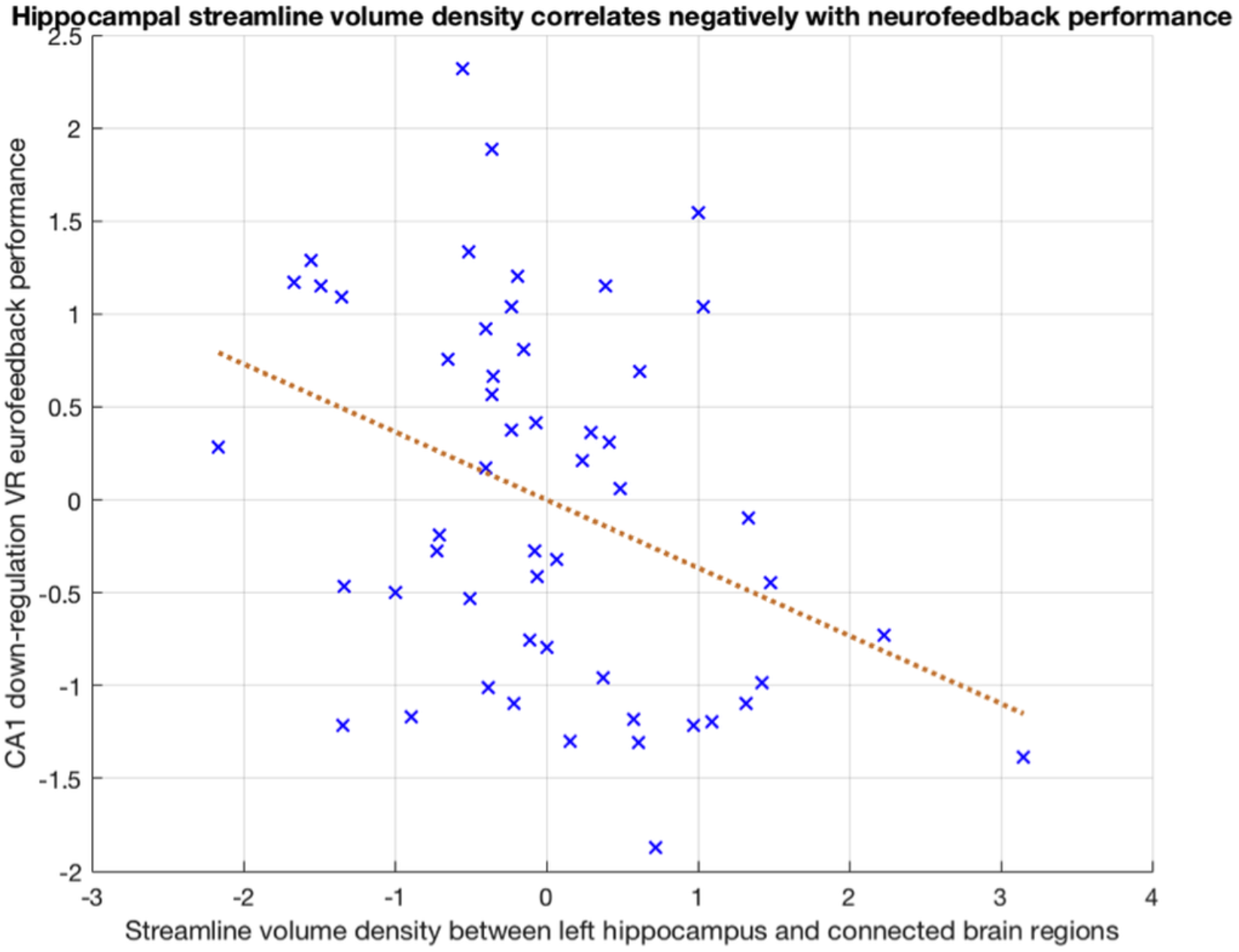
Scatterplot with line of best fit illustrating the correlation between hippocampal streamline density volume and NF performance, corrected for sex, age, APOE genotype, hippocampal volume, cognitive reserve and memory performance, r = −0.371, p = 0.006. Units are in z-scores. For each participant, mean hippocampal streamline density volume was computed across all fibertracts connecting the hippocampus to the five regions that consistently showed bilateral streamlines to the hippocampus, across all 53 participants (i.e. entorhinal cortex, amygdala, thalamus, posterior cingulate and parahippocampal formation).

### Cognitive reserve and brain reserve

Controlling for age, sex, APOE genotype and hippocampal volume, there was no significant correlation between NF performance and cognitive reserve, as measured by the CRQ (r = 0.107, p = 0.444). Similarly, controlling for age, sex, APOE genotype and cognitive reserve, there was no significant correlation between NF performance and brain reserve, as proxied by hippocampal volume normalized by total intracranial volume (r = −0.119, p = 0.394). These measures of cognitive reserve and brain reserve did not correlate either (r = −0.060, p = 0.672).

## Discussion

To our knowledge, this is the first neurofeedback study of the hippocampus, especially in a cognitively healthy population at risk for AD. The results corroborate the established role of APOE genotype as the strongest genetic risk factor for AD [44–45] and suggest that APOE-ε4 alleles impact older adults’ ability to down-regulate hippocampal activation. This interpretation provides a mechanistic explanation with regards to the hippocampal hyperactivity observed in early AD [1] and APOE-ε4 homozygotes [7]. Moreover, the results show that hippocampal down-regulation is not directly affected by other AD risk factors, including cognitive and brain reserve. On the contrary, low SDV, that is related to neurodevelopmental processes [29–30], appears to exert a positive effect on hippocampal down-regulation performance. These results pave the way for novel methods to assess subtle differences in memory function, by proposing neurofeedback performance as a promising metric for preclinical assessments and showing that bilateral hippocampal cohesion during gamified functional neurofeedback, correlates with performance on the most widely used psychometric test for mild cognitive impairment, in cognitively normal participants.

Existing literature provides a feasible explanation for the negative correlation between SDV and NF performance. According to current models, NF learning is enabled via a network comprised of the anterior insular cortex and anterior cingulate cortex, as well as parts of the prefrontal cortex, parietal lobule, ventral striatum and thalamus [45]. It follows that brain regions with direct connections to the NF learning network would be easier to modulate than brain regions with less direct connections. By extension, if a target brain region is connected more heavily to areas that are not part of the NF learning network, it will be more difficult to modulate its activity.

Previous research could not account for the high percentage (up to 30%) of subjects with no improvement in self-regulation performance [46]. Here we have partly addressed this by showing for the first time that variance in NF performance can be accounted by genetic and anatomical factors, providing a foundation for further investigations. Due to the overall negative influence of APOE-ε4 alleles [43–44], good hippocampal down-regulation performance should be tentatively viewed as a resilience factor, rather than as a risk factor. Because brain reserve and cognitive reserve did not exert any strong influence on NF performance, it appears that NF performance may be useful in estimating the independent, functional component of resilience to memory decline, known as neural reserve [8–9].

### Limitations and future directions

Recently we showed that DMN up-regulation and down-regulation learning are negatively correlated [12]. Therefore it is feasible that the present results would be reversed using a hippocampal up-regulation task. It is important to confirm our findings with longitudinal studies investigating both down-regulation and up-regulation, while including sham NF conditions. To this aim, we have developed an upgraded version of the paradigm that enables multi-center investigations within a standardized open-source framework. The paradigm has been designed so it can be easily installed in any hospital or fMRI center with a real-time data setup. It is also compatible with other real-time data acquisition modalities, like EEG, MEG etc. (see section *Code Availability*). Last, given evidence for hippocampal lateralization, here in relation to SDV and previously in relation to functional connectivity with the MCC [3], as well as episodic memory context [11], it is important for future neurofeedback studies of the hippocampus, to investigate the down-regulation of the left and right hippocampi separately.

## Conclusions

We have demonstrated that genetic factors associated to a neurodegenerative disease, can impact self-regulation performance of the brain areas that are most vulnerable to that disease. This opens new avenues for further research applications quantifying and monitoring disease progression in other neurodegenerative diseases, as well as in psychiatric disorders with known genetic risk factors and patterns of functional aberrancy (e.g. schizophrenia, bipolar disorder, etc.). Rt-fMRI VR NF and the implemented open-source experimental paradigm in specific, offer new methods for probing brain function and obtaining subtle metrics that correspond with recognized cognitive and biological processes. These newly developed methods can be used to provide novel metrics of hippocampal function that are based on neuroimaging and are therefore unbiased with regards to the limitations of psychometric questionnaires for measuring memory performance (e.g. self-reporting). Thereby, these methods appear promising for identifying early alterations in brain function due to AD and hold potential for scalable prognostic screening.

## Acknowledgements

This work has received funding from the European Union’s Horizon 2020 research and innovation programme under the Marie Sklodowska-Curie action grant agreement No 707730.

This publication is part of the ALFA (ALzheimer and FAmilies) study. The authors would like to express their most sincere gratitude to the ALFA project participants, without whom this research would have not been possible.

